# miR-137 conferred robustness to the territorial restriction of the neural plate border

**DOI:** 10.1101/2023.09.08.556830

**Authors:** Luciana A. Scatturice, Nicolás Vázquez, Pablo H. Strobl-Mazzulla

## Abstract

The neural plate border (NPB) of vertebrate embryos segregated from the neural plate (NP) and epidermal regions, and comprised an intermingled group of progenitors with multiple fate potential. Recent studies have shown that during the gastrula stage, TFAP2A acts as a pioneer factor in remodeling the epigenetic landscape required to activate components of the NPB induction program. Here we show that *Tfap2a* has two highly conserved binding sites for miR-137 and both display a reciprocal expression pattern at the NPB and NP respectively. In addition, ectopic miR-137 expression reduced TFAP2A, whereas its functional inhibition expanded their territorial distribution overlapping with PAX7. Furthermore, we demonstrated that loss of the *de novo* DNA methyltransferase DNMT3A expanded miR-137 expression to the NPB. Bisulfite sequencing showed a significantly higher level of non-canonical CpH methylation at the promoter of miR-137 when we compared NPB and NP samples. Our finding shows that miR-137 contributes to the robustness of NPB territorial restriction in vertebrate development.

## INTRODUCTION

In all chordates, the ectoderm layer becomes segregated into the neural plate (NP) and the surrounding non-neural ectoderm. The NP subsequently folds or cavitates to form the neural tube, the future central nervous system (CNS), whereas the non-neural ectoderm mostly forms the epidermis of the skin. In vertebrates, at the interface of those two territories resides the neural plate border (NPB). Cells that intermingle at the NPB give rise to four distinct cell lineages: (1) neural progenitors that form the anterior CNS, (2) neural crest cells that form the peripheral nervous system, pigment cells, and much of the bone and cartilage of the face, (3) the cranial placodes that form complex sensory organs such as the inner ear and the olfactory epithelium, and (4) the cranial epidermis (Grocott et al., 2012; Groves and LaBonne, 2014). Particularly, the neural crest and cranial placodes are intimately linked with the evolution of defining characteristics of vertebrates. Whereas basal chordate embryos possess a sharp demarcation between presumptive neural and epidermal fates at this border, much less is known about how cell fates as disparate as neural crest, placode, and neural cells become segregated at the NPB of vertebrate embryos. This segregation may require a tight epigenetic and transcriptional control of both protein-coding genes and non-coding RNAs that contribute to specific genetic programs (Alata Jimenez and Strobl-Mazzulla, 2022; Roellig et al., 2017; Williams et al., 2022a). The biological significance of this region has not only evolutionary relevance but also its implication in several human diseases where genetic or environmental perturbations of these progenitors collectively contribute to an enormous range of birth defects affecting the brain, skull, face, heart, and sensory organs (Butler Tjaden and Trainor, 2013; Gandhi et al., 2020; Pauli et al., 2017; Siismets and Hatch, 2020; Vega-Lopez et al., 2018).

TFAP2A belongs to the TFAP2 family of transcription factors, which includes five paralogous proteins that bind to DNA as dimers (Eckert et al., 2005). This pioneer factor also regulated chromatin accessibility and has been demonstrated to participate in distinct network modules during NPB induction and the later neural crest specification (Rothstein and Simoes-Costa, 2020). Several works using chick embryos demonstrated that TFAP2A is not uniformly expressed in the NPB cells. Doble-immunostaining demarcated TFAP2A expression on the lateral aspect of the NPB, whereas PAX7, another early marker of the NPB (Basch et al., 2006; De Crozé et al., 2011), was mostly detected in the medial border region (Rothstein and Simoes-Costa, 2020). RNA velocity measurements imply that segregation of the NPB begins at the beginning of neurulation (HH6) when the NPB is defined by a discrete subcluster of the ectoderm. However, the NPB also harbors different subclusters heterogeneously expressing several NPB markers including TFAP2A and/or PAX7 (Williams et al., 2022a). This observation reflected the observed segregation of the neural plate border into medial (PAX7+) and lateral (TFAP2A+) regions, as well as highlighting the overlap and combination of genes co-expressed across the NPB. However, the mechanism involved in regulating this complex heterogeneity in the developing NPB is largely unknown.

MicroRNAs (miRNAs) form a group of non-coding RNAs that act in post-transcriptional gene repression and are involved in a myriad of cellular events, including the balance between proliferation and differentiation as well as the spatiotemporal modulation of gene expression during development (Ivey and Srivastava, 2010). Interestingly, miRNAs dramatically expanded in numbers together with the evolution of vertebrate features (Heimberg et al., 2010) and may therefore have contributed to these vertebrate innovations like the formation of the neural crest and placode progenitors residing in the NPB. In this context, miRNAs are well-positioned to function as key regulators of the cross-antagonism to refine the transcriptional outcome and modulate lineage specification at the NPB. Furthermore, the regulatory elements of those miRNAs may function as integrated transcription factor binding platforms, where environmental signaling cues are interpreted in a context-dependent manner (Buecker and Wysocka, 2012).

Here, we explore the hypothesis that miRNAs could have a role in the spatial control of TFAP2A expression and robustness to the process of NPB cell specification. We identified miR-137 as a direct regulator, characterized their tissue distribution, and performed gain– and loss-of-function experiments demonstrating their role in NPB definition. Next, demonstrated the territorial restriction of miR-137 expression exerted by the DNA methyltransferase 3A and identified differential non-canonical CpH methylation at their promoter region in NPB progenitors. Finally, we demonstrated a feedback loop regulation between miR-137 and TFAP2Aa.

## RESULTS

### MiR-137 as a possible post-transcriptional regulator of TFAP2A

Given that *Tfap2a* is a pioneer factor regulating NPB specification, we performed an *in silico* analysis to identify candidate miRNAs that might regulate their territorial restriction. Through this, we found that miR-137 is the only one having two binding sites with a higher probability of preferential conservation across vertebrates in the 3’UTR of *Tfap2a* mRNAs **(Fig. 1A-S1)**. One of the sites is the only one having the highest base complementarity (8mer) between the mRNA and the seed region. Moreover, from all the miRNAs targeting *Tfap2a*, miR-137 has the highest probability of preferentially conserved targeting (aggregate P_CT_=98) **(Table S1)**. MiR-137 is a brain-enriched miRNA that plays an important role in regulating embryonic neural stem cell fate determination, neuronal proliferation and differentiation, and synaptic maturation. Its dysregulation results in alterations in the gene expression regulation network of the nervous system, which in turn, induces mental disorders in humans, including schizophrenia susceptibility (Yin et al., 2014).

**Figure 1:**
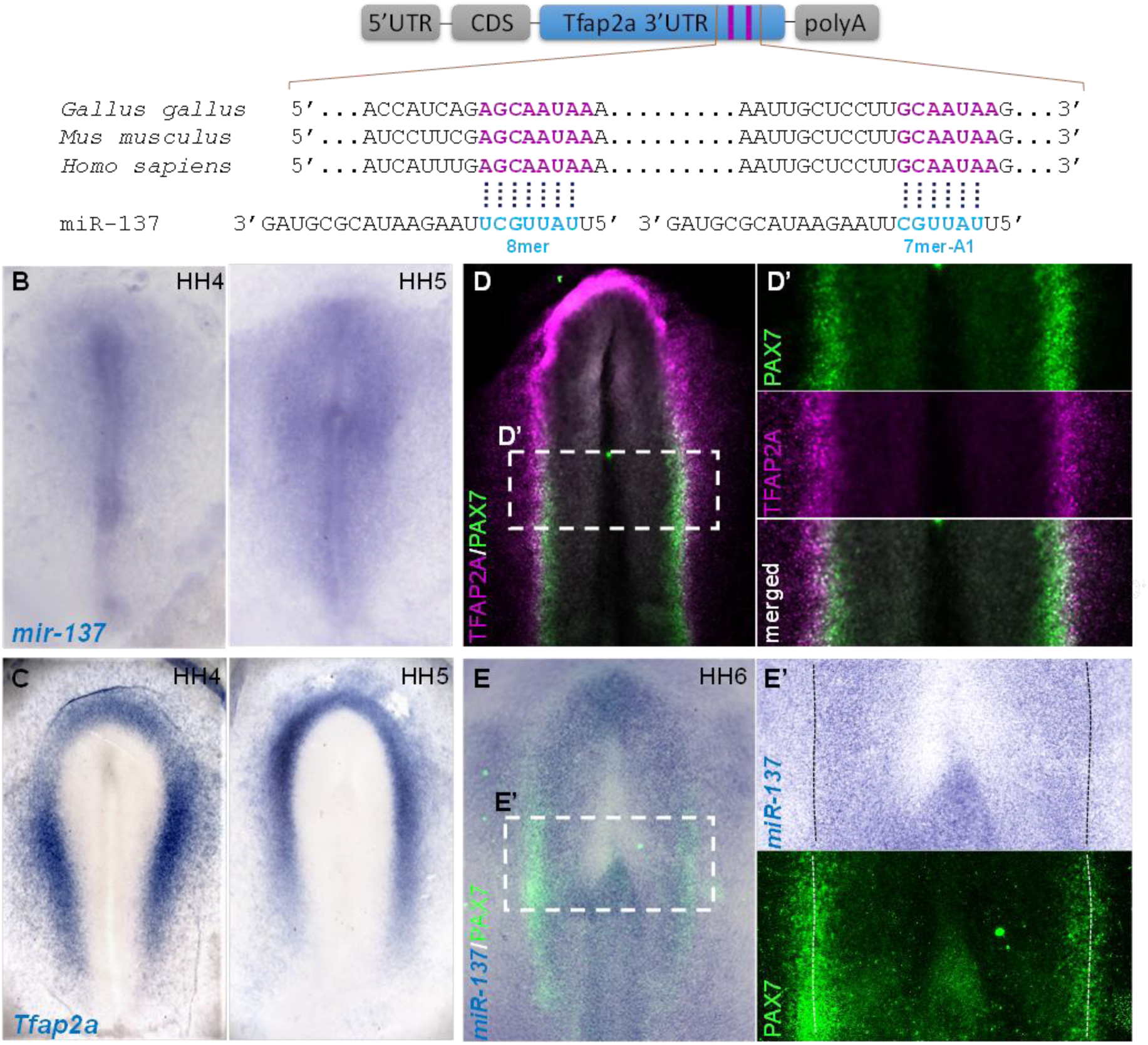
miR-137 expression is opposite to the putative target gene Tfap2a. (**A**) Mature miR-137 has two binding sites on the 3’UTR of Tfap2a mRNA which are highly conserved in vertebrates. **(B)** Whole-mount *in situ* hybridization using LNA-DIG-labeled probes reveals specific expression of mature miR-137 in the neural plate at HH4-5, which is opposite to Tfap2a **(C)** expression detected in the neural plate border and non-neural ectoderm. **(D)** Immunostaining evidencing that the neural plate border expressed TFAP2A (magenta) on the medio-lateral side, and PAX7 (green) on the medio-central side. **(D’)** Magnification at the level shown in D. **(E)** MiR-137 in situ hybridization followed by PAX7 (green) immunostaining demonstrating the limits of miR-137 in the middle of PAX7 expression (dotted line E’).

Once the candidate was identified, we wanted to examine the expression pattern of miR-137 at different stages in early chick embryos and compare it with the pattern of *Tfap2a* mRNA. MiRNAs and their targets have been reported to be often mutually exclusive expression in tissues during embryonic development, especially in neighboring tissues derived from common progenitors (Ebert and Sharp, 2012). In this sense, miRNAs may act to reinforce the transcriptional gene expression program by suppressing random fluctuations in transcript copy number at territorial boundaries, thus allowing the generation of sharper limits in the expression of their targets. As a first step, we examined the expression pattern of the mature miR-137 transcripts using *in situ* hybridization in early chick embryos. By utilizing a locked nucleic acid (LNA) digoxigenin-labeled probe we found that miR-137 was first detected at gastrula stages (HH4-5. Hamburger & Hamilton stages) on the neural plate **(Fig. 1B-S2)**. Interestingly, *Tfap2a* transcript, which is highly detected in the NPB and non-neural ectoderm, seems to have an anti-correlated expression with respect to miR-137 **(Fig. 1C)**. This becomes very evident when miR-137 transcript is co-immunostained with the NPB marker PAX7 **(Fig. 1D-E)**. This evidence is consistent with the intriguing possibility that miR-137 may have some functional role in the regulation of *Tfap2a* spatial distribution.

### Ectopic miR-137 causes TFAP2A reduction at the NPB

In order to study the *in vivo* effect of ectopic miR-137 expression on *Tfap2a* transcript at the NPB we designed an overexpressing vector containing the pri-miR-137 under the control of CAG promoter (pOmir-137). This vector, or the empty version (pOmir), was unilaterally co-injected with a GFP reporter plasmid into HH4 embryos, followed by electroporation and allowed them to develop until HH7 **(Fig. 2A)**. Prior to functional analysis, we demonstrated by stem-loop-RT-qPCR that this vector is indeed able to overexpress miR-137, comparing the injected side (IS) with the uninjected side (UIS) of the same group of electroporated embryos **(Fig. 2B)**. To visualize the effect of ectopic miR-137, we evaluated by ISH and IHC the *Tfap2a* expression. Our results demonstrated that both *Tfap2a* mRNA and protein were reduced at the pOmir-137 injected side compared with the contralateral side, or embryos injected with the control vector pOmir **(Fig. 2C-D)**. Taken together, our results highlight the possible regulatory role of miR-137 in the *Tfap2a* territorial restriction.

**Figure 2:**
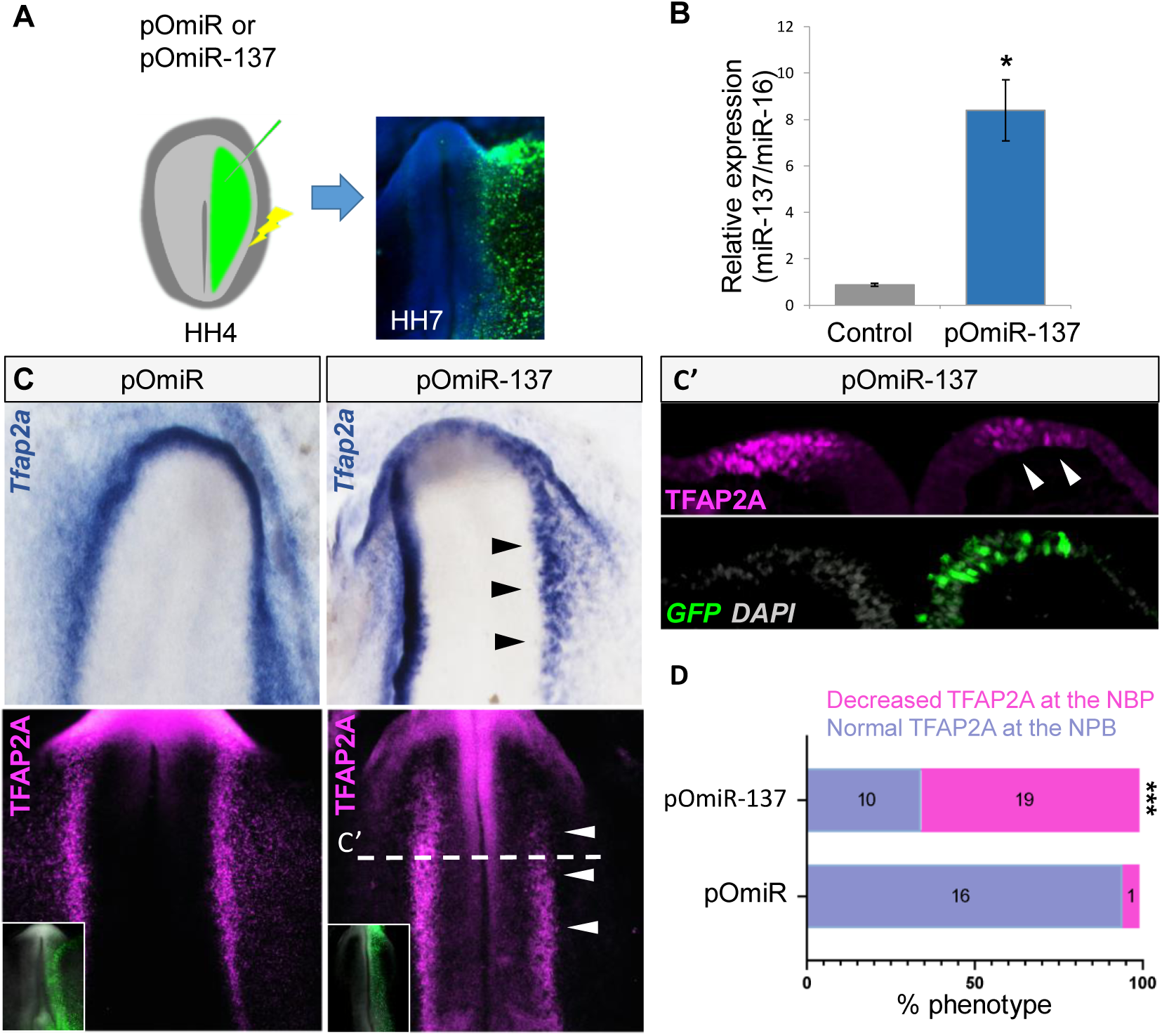
miR-137 overexpression reduced TFAP2A expression. (**A**) Schematic diagram of the overexpressing (pOmiR-137) or empty control (pOmiR) vectors injected and electroporated in the right side of the embryos at gastrula stage (HH4) and cultured until mid-neurula stage (HH7). **(B)** Electroporation of pOmiR-137 vector successfully overexpress a mature *miR-137* evidenced by stem-loop-RTqPCR compared with the control uninjected side of the same group of embryos (n=3). Asterisk (*) indicates significant differences (*P*<0.05) by Student’s *t*-test. Values are means ± SEM. **(C)** *In situ* hybridization and immunostaining for TFAP2A show their reduction in the pOmiR-137 injected side compared with the uninjected side of the same embryos and with embryos injected with the control pOmiR vector. **(C′)** Transverse section at the levels shown in C. **(D)** The percentages of embryos showing a phenotype (decreased *Tfap2a* expression at the neural plate border) on the side injected with pOmiR-137 and pOmiR. Numbers in the graphs represent the numbers of analyzed embryos. Asterisk (***) indicates significant differences (*P*<0.001) by contingency table followed by χ2 test.

### Loss of miR-137 function leads to TFAP2A invasion of the NP territory

Given that miR-137 is expressed in the NP territory and its ectopic expression in the NPB causes repression in TFAP2A expression, we next asked whether their loss of miR-137 function in the NP would expand the TFAP2A territory to this region. To test this possibility, we utilized two approaches. First, generating a “sponge” vector containing 10 repeated miR-137 antisense sequences (pSmiR-137) to sequester endogenous miR-137(Kluiver et al., 2012). We have also designed a control sponge vector containing a scramble miR-137 sequence (pSmiR-scra). Second, we used an IDT inhibitor (Ihn-miR137) designed to be perfectly complementary to the mature miRNA-137 sequence, producing a very stable duplex that prevents the miRNA from binding to its intended targets and is capable of even displacing the natural passenger strand in double-stranded miRNA (Lennox et al., 2013). As a control, we used a non-targeting control (Ihn-NC5). In both cases, the sponge plasmids or the miRNA inhibitors were transfected into the right side of the embryos as we mentioned before. The results utilizing the sponge vector demonstrated the expansion of *Tfap2a* transcripts toward the NP territory, as well as we observed a more diffuse border compared to the uninjected side or control embryos injected with pSmiR-scra **(Fig. 3A)**. It has been previously reported that TFAP2A demarcates de lateral aspect of the NPB extending toward the non-neural ectoderm, and on the other side PAX7, another well-known NPB marker, is mostly enriched in the medial border region of the NPB (Williams et al., 2022a). This transcriptional heterogeneity of the progenitors located in the NPB seems to be very important for the neural crest and placode specification. Based on our hypothesis, miRNA-137 may act to suppress random fluctuations in TFAP2A expression generating a sharper delimitation. To evaluate this, we manipulated miR-137 expression *in vivo* by transfecting half embryos with ihn-miR137 and determined the effect on TFAP2A and PAX7 protein distribution by immunofluorescence. The results show an expansion of TFAP2A expression over the PAX7 territory in the ihn-miR137 injected side compared to the control side **(Fig. 3B-E)**. These analyses demonstrated that TFAP2A expression is expanded to more medial territories, overlapping with PAX7 distribution, in ihn-miR137 transfected side (the negative control inh-NC5 displayed no effect) **(Fig. 3D)**. To further evaluate these results, we performed transversal sections in the anterior zone of both miR-137 inhibitor-treated and control embryos.

**Figure 3:**
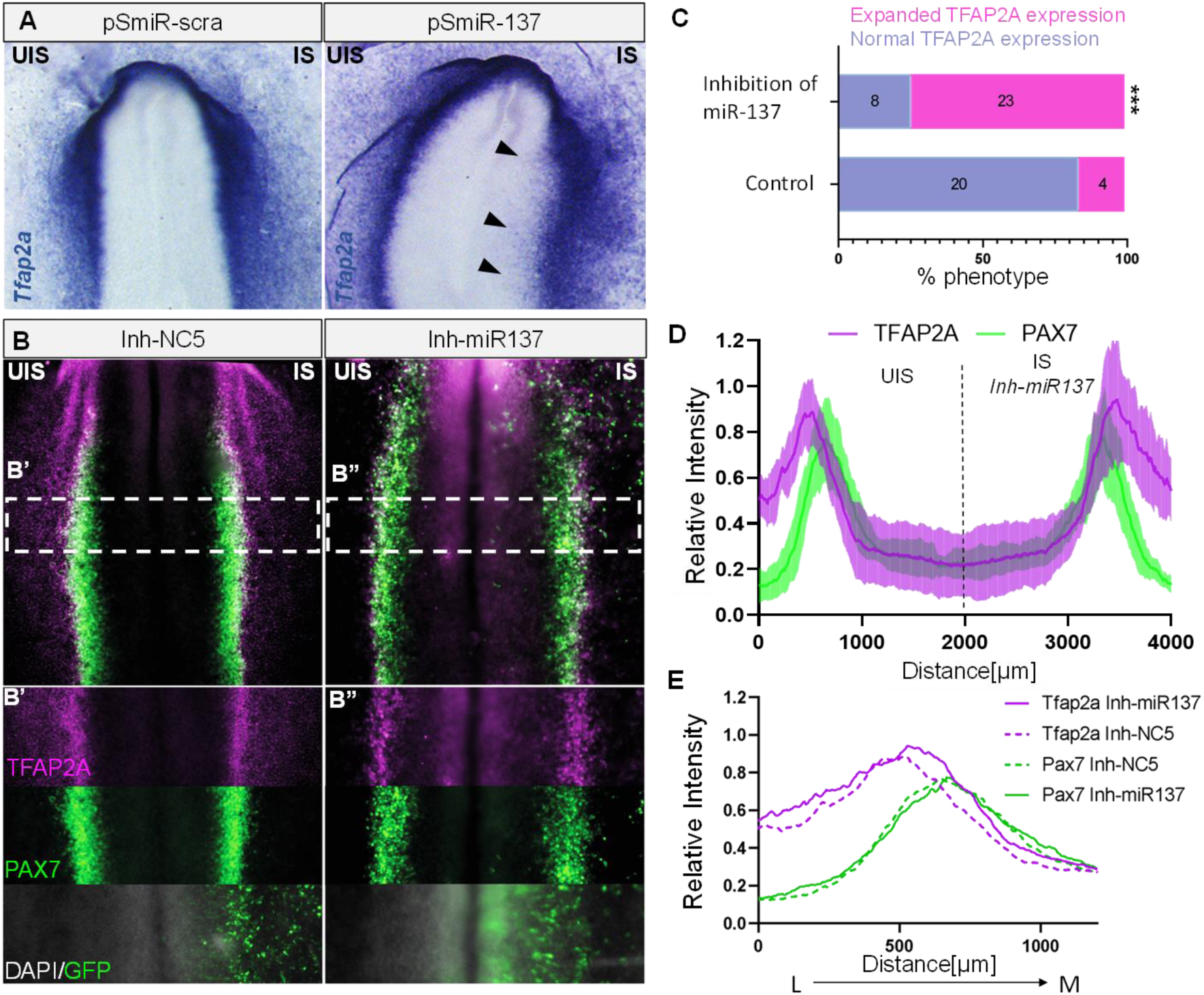
Loss of miR-137 causes Tfap2a territorial expansion. (**A**) Embryos electroporated with *miR-137* sponge vector (pSmiR-137) presented *Tfap2a* expansion to the neural plate territory (black arrowhead), as shown by *in situ* hybridization compared with the uninjected side of the same embryos (Left side) or injected with a scramble sponge vector (pSmiR-scra). **(B)** Co-immunostaining for PAX7 (green) and TFAP2A (magenta) after injection with miR-137 inhibitor (Inh-miR137) demonstrated an expansion of TFAP2A+ territory over PAX7+ cells compared with the uninjected side of the same embryos or injected with a non-targeting control inhibitor (Inh-NC5). **(B′, B”)** Independent channels at the levels shown in B. **(C)** The percentages of embryos showing a phenotype (expanded *Tfap2a* expression) on the side injected with Inh-miR137 and Control. Numbers in the graphs represent the numbers of analyzed embryos. Asterisk (***) indicates significant differences (*P*<0.001) by contingency table followed by χ2 test. **(D)** Relative intensity profiles of PAX7 (green) and TFAP2A (magenta) protein expression across the embryos (n= 7) demonstrating their overlapping distribution on the injected side with Inh-mR137 (right side). Dark lines indicate average across all embryos, with standard deviation indicated by shaded regions. **(E)** Line traces of average relative intensities for TFAP2A and PAX7 expression comparing injected sides with inh-miR137 (solid lines) and control inh-NC5 (dotted lines) across the neural plate border from lateral to medial. UIS, injected side; IS, injected side, L, lateral, M, medial.

Interestingly, we found that the TFAP2A expression is expanded toward the medial NPB, overlapping with PAX7 expression on the Inh-miR137 injected side compared to the uninjected side of the same embryos and the Inh-NC5 injected side (**Fig. 4A**). These results indicated that miR-137 is required for proper *Tfap2a* territorial restriction during NPB definition. The induction and specification of different cell lineages of the NPB (neural crest cells in the medial zone and placode cells in the lateral zone) is a consequence of the heterogeneous distribution of TFs along the territory that generates a precise transcriptional context for each precursor. Changes in the signaling gradients could affect their progenitors, thus leading to anomalous induction and differentiation. With this in mind, we evaluated whether the number of cells along the NPB expressed TFAP2A and PAX7 deferentially upon miR-137 inhibition. Using the FIJI tool and the images obtained from the transversal sections, we counted cells that only express TFAP2A (cells only TFAP2A+), PAX7 (cells only PAX7+), and those co-expressing both proteins (cells PAX7+/TFAP2A+). Indeed, we found that in the embryos treated with the miR137 inhibitor, there are significantly more cells co-expressing both markers compared to the control (**Fig. 4B**). Therefore, our result suggests miR-137 could be regulating the expression of TFAP2A in the most medial NPB cell population, at the edge with the NP, for the correct induction of the progenitors of this region and the specification of their derivatives, particularly the neural crest cells.

**Figure 4:**
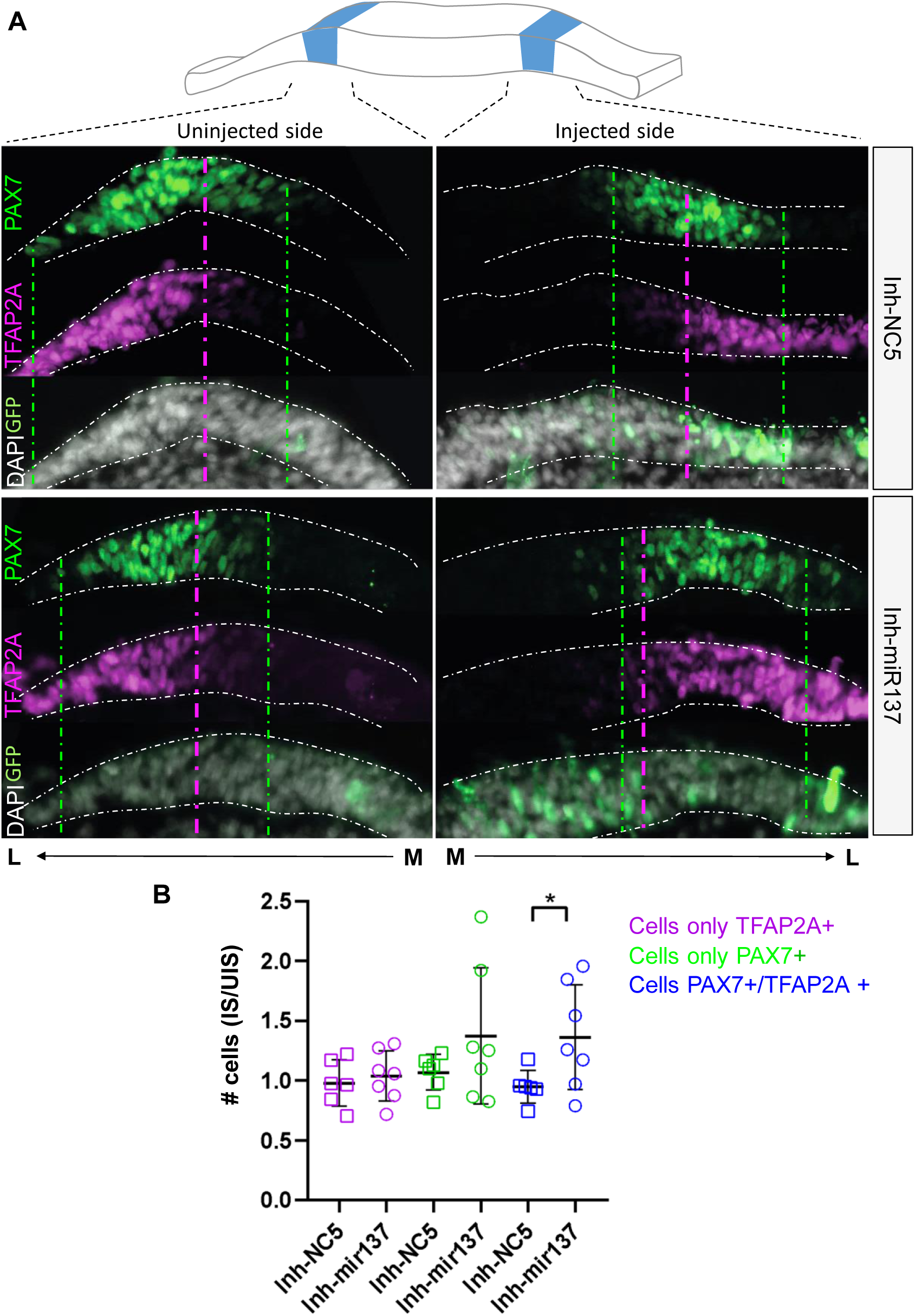
Loss of miR-137 increased TFAP2A+/PAX7+ co-expression in single cells. (**A**) Transverse sections on HH7 embryos immunostained for PAX7 (green) and TFAP2A (magenta) showing the uninjected (UIS) and injected side (IS) with Inh-miR137 or Inh-NC5. M, medial; L, lateral. Dotted lines delimited the PAX7 and TFAP2A territories **(B)** Scatterplot quantitation of cells ratios (total # of counted cells at IS/UIS) for only PAX7+ (green), only TFAP2A+ (magenta), or both PAX7+/TFAP2A+ (blue) on Inh-NC5 (n=6) and Inh-miR137 (n=7) treated embryos. Asterisks indicate significance (*P*<0.05) calculated using a Student’s *t*-test. Values are means ± SD.

### DNMT3A is required for miR-137 territorial restriction

MiRNAs are often embedded in CpG islands and are susceptible to DNA methylation as a major mechanism to repress their expression (Shukla et al., 2020; Wiklund et al., 2010), and miR-137 is not an exception (Mahmoudi and Cairns, 2017). Earlier studies conducted by our group have shown that the *de novo* DNA methyltransferase DNMT3A plays a crucial role in neural versus neural crest specification. Particularly, DNMT3A was found to be highly expressed in the NPB during gastrulation (Hu et al., 2012), thereby making it an ideal candidate for repressing miR-137 expression in this region. To examine this possibility, we transfected half embryos with a previously characterized morpholino known to block the translation of DNMT3A (DNMT3A-MO). Subsequently, we performed ISH to visualize the expression of miR-137. The results show that depletion of DNMT3A resulted in an expanded miR-137 expression to the NPB territory compared with the uninjected side or embryos treated with Control-MO **(Fig. 5A)**. This result suggests that DNA methylation may be involved in the territorial restriction of miR-137 expression.

**Figure 5:**
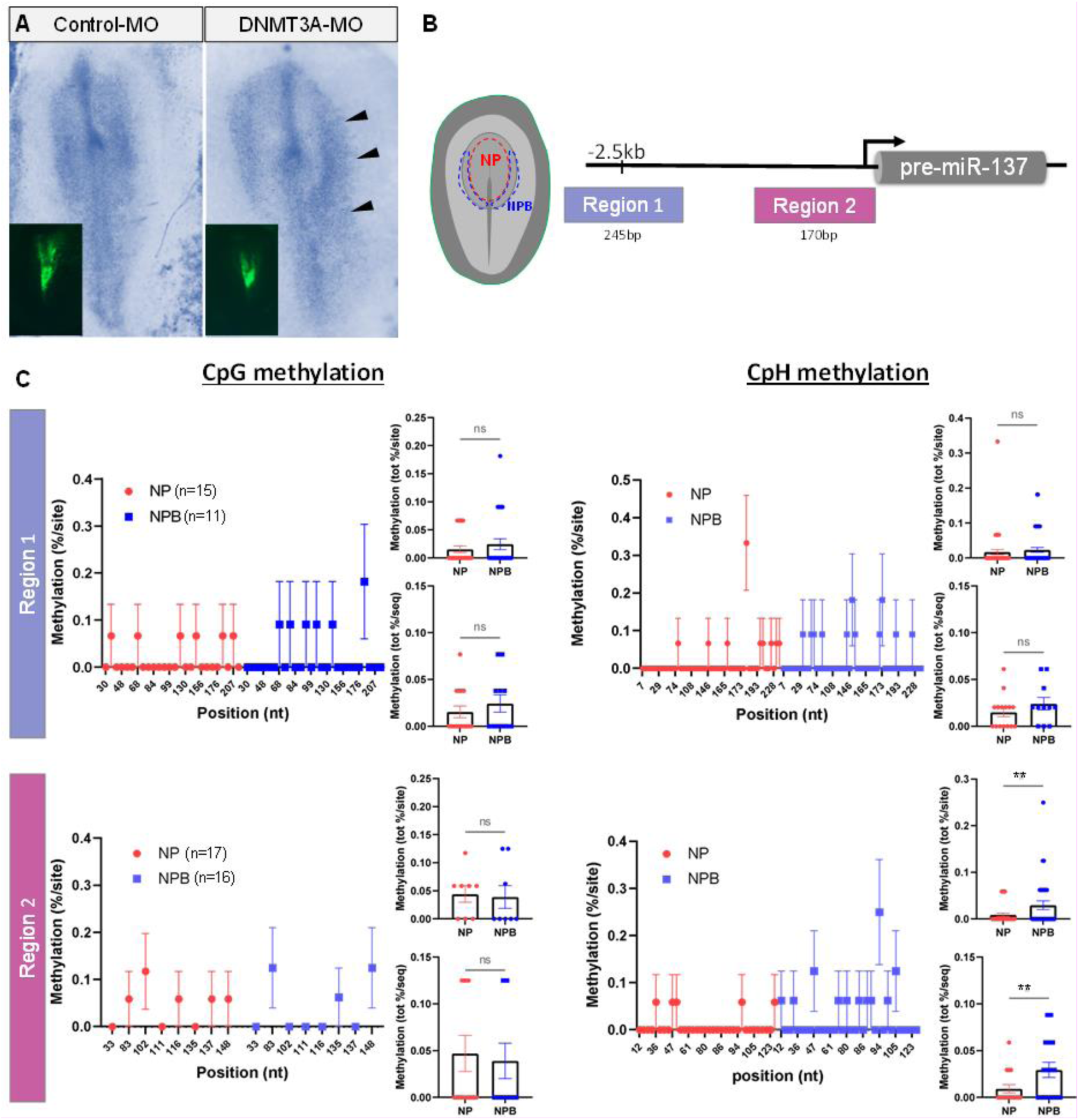
Loss of DNMT2A expands miR137 expression to the neural plate border where its locus is normally highly methylated. (**A**) Morpholino-mediated loss of DNMT3A (DNMTA-MO) results in an expanded expression of *miR-137* into the neural plate border (black arrowheads) evidenced by whole-mount *in situ* hybridization on the injected side compared with the uninjected side of the same embryos and injected with Control-MO. **(B)** A tissue dissection scheme involving the neural plate (NP, red) and neural plate border (NPB, blue) was employed to investigate the methylation patterns of two regions (Region 1 and Region 2) near the miR-137 locus using bisulfite sequencing. **(C)** Percentage of methylation on CpG and CpH (A, T or G) sites located in Region 1 and Region 2 from neural plate (NP, red dots) and neural plate border (NPB, blue squares) samples. The percentages of individual methylated sites, total methylation per site, and total methylation per sequence are shown for CpG and CpH in each region. We found a significantly higher percentage of methylated CpHs in Region 2 of the NPB compared with NP samples. The numbers in brackets indicate the number of analyzed sequences. Asterisks indicate significance (*P*<0.05) as calculated using a Student’s *t*-test. Values are means ± SEM. ns, non-significant differences.

### The miR-137 locus is differentially methylated among NP and NPB territories

Considering the above results, we asked whether there was in fact a differential methylation between NPB and NP territories in two regulatory regions of miR-137 locus **(Fig. 5B)**. These regions were selected, one near the pre-miR-137 (Region 2), and the other at the putative proximal promoter (Region 1) previously described to be regulated by DNA methylation (Szulwach et al., 2011). We dissected the NP and NPB from HH5+ embryos and analyzed them by using bisulfite conversion to visualize the canonical (CpG) and non-canonical (CpH) methylations on those specific regions. Our analysis of Region 1 revealed no significant differences in the percentage of CpG or CpH methylations when examining the region as a whole (% of methylation/sequence) or at individual sites (% of methylation/site) **(Fig. 5C)**. Interestingly, when we analyzed the Region 2 we observed a significant increase in the percentage of methylation per sequence and sites in CpH, while there was no corresponding increase in CpG. Based on our findings, it appears that the higher level of non-canonical methylation in the NPB, as compared to the NP, could be responsible for the territorial restriction of miR-137.

## DISCUSSION

There is increasing evidence about the role of microRNAs during embryonic development in reinforcing the transcriptional gene expression program at territorial boundaries, acting as fine-tuning regulators, and generating more pronounced boundaries in the expression of key transcription factors for proper territorial delimitation (Hornstein and Shomron, 2006). The NPB cannot be defined solely by a specific set of genes uniformly and/or exclusively expressed in all their cells. Instead, its distinct multipotency arises from a combination of genes differentially expressed on them. Therefore, we propose adding miRNAs as new participants in Roellig’s binary competition model (Roellig et al., 2017), wherein territory-specific transcription factors exhibit mutual exclusivity. Thus, miRNAs may act as fine regulators, limiting fluctuations in the number of transcripts, resulting in more precise territorial transitions and conferring robustness to the NPB cell specification process. Particularly, our study highlights the key role of a single microRNA, miR-137, in the territorial restriction of the NPB master transcription factor, *Tfap2a*. Our results show that ectopic expression of miR-137 in the NPB territory reduces both *Tfap2a* mRNA and protein; whereas loss of miR-137 function in the neural territory generates its expansion over the medial NPB, overlapping with PAX7 (see hypothetical model in **Fig. 6**). The transition from a pluripotent to a specialized cell state involves successive stages with characteristic transcriptional states. One of these states, known as “*primed”* state, is characterized by the co-expression of transcription factors from different cell lineages. This is the case described in cells comprising the NPB, where a subset of cells co-expressing different TFs gives plasticity to these progenitors, able to respond to several signals and contribute to different cell lineages that arise from this territory (Hu et al., 1997; Laslo et al., 2006; Olsson et al., 2016). However, what segregated those *primed* progenitors to cell lineage commitment and specification toward specific lineages? In this regard, our results show that repression of miR-137 increases the number of precursors co-expressing TFAP2A and PAX7, which could influence the proportion of progenitors in *primed* states and, consequently, influence the cell populations that derive from them. Although our analysis is limited by the use of only two markers, it reinforces the idea that miR-137 limits TFAP2A expression in the medial NPB by decreasing its transcriptional random fluctuations at the boundaries.

**Figure 6:**
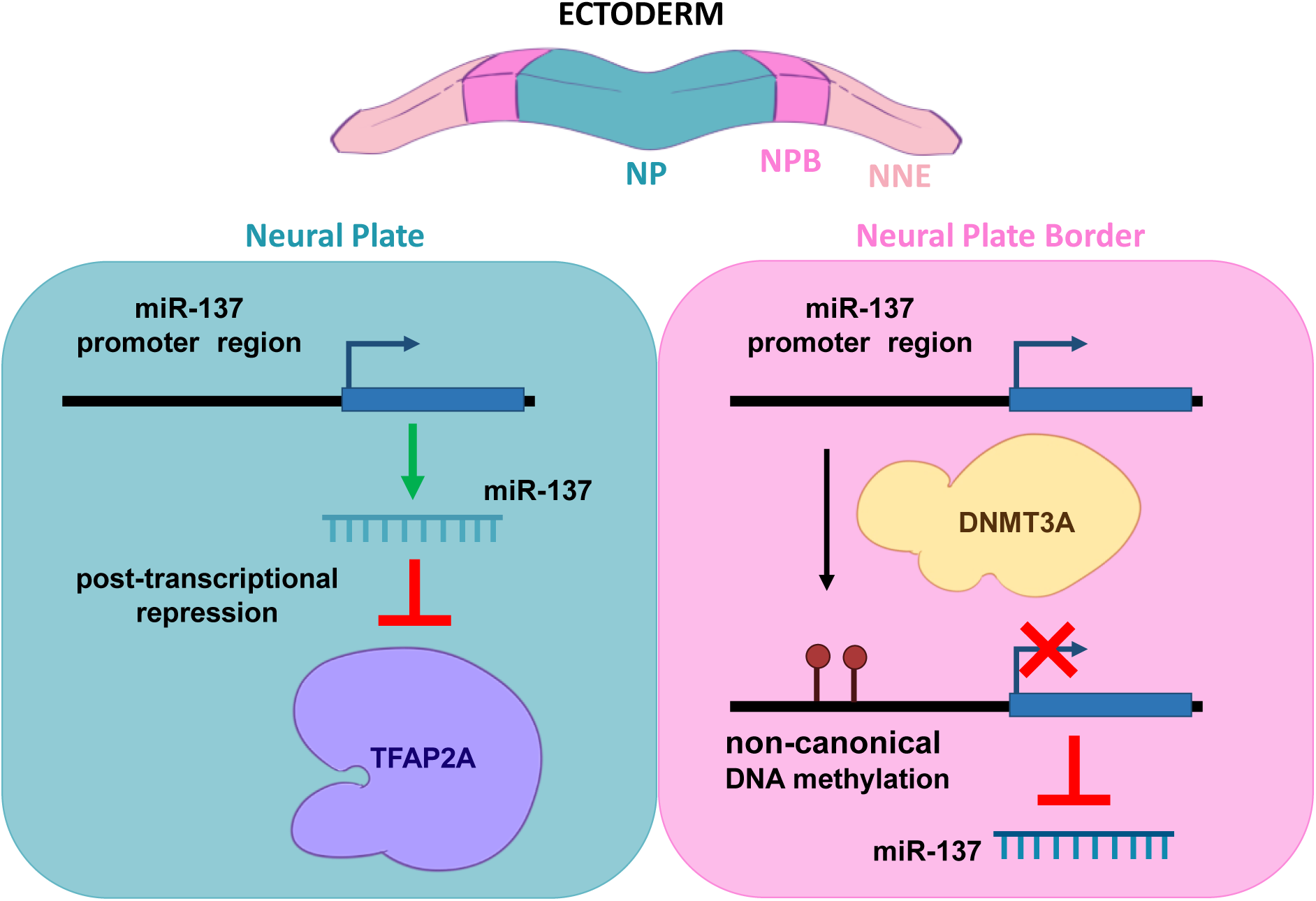
Hypothetical model. Our results show that, during NPB specification, miR-137 is expressed in the NP and limits the expression of TFAP2A in this territory. On the other hand, miR-137 expression is restricted by the action of DNMT3A in the NPB via non-canonical methylation of its promoter region.

Epigenetic regulation by DNA methylation has been described for several miRNAs, including miR-137 (Balaguer et al., 2010; Kozaki et al., 2008; Langevin et al., 2010; Sańchez-Vaśquez et al., 2019; Szulwach et al., 2011). Specifically, hypermethylation of miRNA promoters, mostly embedded in CpG islands, leads to their silencing in different cell types (Balaguer et al., 2010; Chen et al., 2011; Kozaki et al., 2008; Langevin et al., 2010). Particularly, methylation in the regulatory regions of miR-137 has been reported to repress its expression playing a key role in modulating adult mouse neurogenesis (Szulwach et al., 2010). Here, we found that the miR-137 promoter region has a higher level of non-canonical methylation (CpH) in the NPB territory compared with the NP territory, and the lack of DNMT3A expanded the miR137 expression to the NPB territory. Given that DNMT3A exhibits a specific preference for methylating asymmetric CpH sites and is primarily expressed in the NPB during embryonic development (Hu et al., 2012; Jang et al., 2017), we propose its involvement in the transcriptional repression of miR-137 in this region through a non-canonical methylation mechanism acting on its promoter. Nevertheless, despite the universal occurrence of non-CpG methylation, its function, and underlying mechanisms remain elusive and subject to controversy (Patil et al., 2014; Smith and Meissner, 2013). There are different perspectives among researchers regarding non-CpG methylation. Some believe it is a by-product resulting from the hyperactivity of non-specific *de novo* methylation of CpG sites (Smith and Meissner, 2013; Ziller et al., 2011). Conversely, others argue that non-CpG methylation is associated with gene expression and tissue specificity (Barrès et al., 2009; Lister et al., 2011; Ma et al., 2014). Based on this, recent studies have revealed that non-CpG methylation has been found to be enriched in various cell types, including ES cells (Laurent et al., 2010; Lister et al., 2009), induced pluripotent stem cells (iPS cells) (Lister et al., 2011; Ma et al., 2014), oocytes (Guo et al., 2014; Tomizawa et al., 2011), neurons and glial cells (Lister et al., 2013), although it is rare in most differentiated cell types. Considering this, in our study it appears that the NPB exhibits higher levels of non-CpG methylation compared to the NP. This observation may indicate that the NPB is maintained in a more *naïve* stage of differentiation, as suggested by various authors in previous studies (Roellig et al., 2017; Williams et al., 2022b). However, further studies are needed to unambiguously demonstrate this.

At the mechanistic level, miRNA function is intricately linked with various gene regulatory processes, notably mRNA transcription, splicing, and stability, which collectively enhance the system’s robustness. A persistent inquiry within the field, applicable not only to neural plate border development, revolves around the extent to which miRNAs function as ‘switches’ compared to being ‘fine-tuners’ of gene expression. Importantly, the modus operandi of miRNAs is dictated by spatiotemporal context while exhibiting a preference for targeting genes with low expression levels, leading to the most significant reduction in noise (Ebert and Sharp, 2012; Paulsson, 2004). Moreover, the phenomenon of combinatorial miRNA regulation (Shukla et al., 2020) further enhances overall noise reduction by providing strong repression to endogenous genes while incurring minimal additional noise from miRNA pools. Consequently, we postulate that miRNA regulation emerges as a potent mechanism to reinforce cellular identity by mitigating undesirable gene expression fluctuations within the neural plate border progenitors.

## MATERIALS AND METHODS

### Embryos

Fertilized White Leghorn chick eggs were purchased from “Escuela de Educación Secundaria Agraria de Chascomús” and incubated at 38°C until embryos reached the desired stage. Chicken embryos were collected and staged according to the criteria of Hamburger and Hamilton (Hamburger and Hamilton, 1992). For in situ hybridization, embryos were fixed overnight at 4°C in PBS-0.1% tween (PBS-Tw) (pH 7.4) containing 4% paraformaldehyde (PFA), dehydrated, and stored in methanol. For immunochemistry, embryos were fixed for 15 min at room temperature in PBS-0.5% Triton (PBS-T) containing 4% PFA, and processed immediately.

### Electroporation of morpholino, vectors, and inhibitors

Embryos were electroporated at stage 3-4 as previously described (Sauka-Spengler and Barembaum, 2008). We used a previously tested antisense morpholino DNMT3A-MO and Control-MO both at 0.75mM (Hu et al., 2012). For the miR137 over-expression experiments, 2 μg/μl of pOmiR-137 or pOmiR vector were injected. For the loss of the miR-137 function, we generated specific and scrambled “sponge” vectors (pSmiR-137 and pSmiR-scra) as previously described (Kluiver et al., 2012). Alternatively, we used the commercial IDT inhibitors (Ihn-miR137 and Ihn-NC5 as non-targeting control) diluted at 10µM in 10 mM Tris pH 8.0 (Lennox et al., 2013). Vectors, morpholinos, and IDT inhibitors were all co-injected with 1µg/µl of carrier DNA. The injections were performed by air pressure using a glass micropipette to target half-side embryos, leaving the opposite side as control. Electroporation was made with five 50 ms pulses of 5.2 V, with intervals of 100 ms between each pulse. Embryos were cultured in 0.75 mL of albumen in tissue culture dishes and incubated at 38°C until the desired stages. All embryos were screened prior to further analysis; embryos with weak and/or patchy electroporation or with strong morphological abnormalities were discarded. All the utilized primers, oligos, morpholinos, and inhibitors are listed in **Table S2**.

### RNA preparation and Stem-loop-RT-qPCR

RNA was prepared from individual chick embryos using the isolation kit RNAqueous-Micro (Ambion) following the manufacturer’s instructions. The RNA was treated with amplification grade DNaseI (Invitrogen) and then reverse transcribed to cDNA using a reverse transcription kit (SuperScript II; Invitrogen) with Stem-Loop-miRNA primers (see Table S2) as previously described (Chen et al., 2005). As normalization control we used miR-16, which was previously validated by our group (Sańchez-Vaśquez et al., 2019). QPCRs were performed using a 96-well plate qPCR machine (Stratagen) with SYBR green with ROX (Roche).

### *In situ* hybridization (ISH)

Whole-mount chick ISH for mRNAs and for microRNA was performed as described previously (Acloque et al., 2008; Darnell et al., 2006). Antisense LNA probes for miR-137 (see Table S2) used in the assay were obtained from Exiqon. Digoxigenin-labeled probes were synthesized from linearized vectors containing full-length cDNAs of *Tfap2a*. Hybridized probes were detected using an alkaline phosphatase-conjugated anti-digoxigenin antibody (Roche, 1:200) in the presence of NBT/BCIP substrate (Roche). Embryos were photographed as a whole-mount using a ZEISS SteREO Discovery V20 Stereomicroscope (Axiocam 512 color) and Carl ZEISS ZEN2 (blue edition) software.

### Cryosectioning

For histological analysis, embryos were incubated in 5% sucrose (in PBS) for 2h at room temperature and subsequently transferred to 15% sucrose and incubated overnight at 4°C. After that, embryos were transferred and incubated in 7.5% gelatin in 15% sucrose for 4 h at 37°C, then, they were frozen with liquid nitrogen and immediately stored at –80°C for cryosectioning. Transverse sections of 10-15µm were generated at the Histotechnical Service (INTECH. Lic. Gabriela Carina Lopez) and used for immunostaining.

### Immunohistochemistry

Embryos or sections were washed in PBS with 0.5% Triton (PBS-T) and subsequently blocked with 5% FBS in PBS-T for 3h at RT. Embryos or sections were incubated in primary antibody solution at 4°C overnight. Primary antibody used: mouse monoclonal anti-TFAP2A IgG2b (Developmental Studies Hybridoma Bank, 1:50) and anti-PAX7 IgG1 (Developmental Studies Hybridoma Bank, 1:10). The secondary antibodies used were goat anti-mouse IgG2b 594 and goat anti-mouse IgG1 647 (all from Molecular Probes, 1:500). After several washes in PBS-T, embryos and sections were mounted and imaged by using Carl ZEISS Axio observer 7 inverted microscope (Axio observer Colibri 7, Axiocam 305 color, Axiocam 503 mono) and Carl ZEISS ZEN2 (blue edition) software.

### Quantification

Fiji software (Schindelin et al., 2012) was used to measure the intensity of protein expression on Zeiss.czi files. As described by Roellig *et al* (Roellig et al., 2017). Using the segmented line tool, a line of about 215 microns in width and 4 mm long was drawn across whole embryos from the uninjected side (UIS) to the injected side (IS). The intensity was measured as grey values. Background and a reference area were measured by placing a fixed-sized oval (1087 square pixels) (reference areas: for PAX7 highest intensity area in neural plate border/neural fold, and TFAP2A epidermis on lateral side). The background was subtracted from marker expression intensity and the intensity of the reference area. Negative numbers were changed to zero. The PAX7 intensity curve was used as a reference to align the curves obtained from each embryo.

For cell number quantitation, Fiji software was used to analyze the cross-section images. Using the segmented line tool, a line of about 215 microns width was drawn, and the ectoderm was used to define the area of interest (from the medial to the lateral side of the ectoderm). For each channel (DAPI, PAX7 y TFAP2A): the tissue background was removed by placing a fixed-sized oval (421 square pixels). Images were linearized using the Straighten tool. Rolling ball tool (Radius=50 pixel) was used to subtract possible residual, and Median (Radius=2.0 pixel) and Gaussean Blur (Sigma radius=2.0 pixel) filters were used to improve the tonality of the image. Through Find Max (Prominence=10) a segmented particle mask was obtained with each peak representing a cell. A second mask, the Threshold tool (Huang), was used to obtain the total area where each marker is located (small particles were eliminated by Analyze Particles). The total number of cells expressing one marker was obtained by multiplying the DAPI segmentation particles mask and the area mask of each marker, as well as marker overlap in the case of cells co-expressing PAX7+ and TFAP2A+. Data were transferred to Microsoft Excel, values of each embryo (4 sections/embryo, n = 7) were averaged, then the ratio between injected side/uninjected side was obtained for each set of cells (PAX7, TFAP2A, and overlapping), and the significance was calculated using Student’s *t*-test using Prism 9 GraphPad.

### Bisulfite sequencing

Samples were obtained by micro-dissecting embryos at HH5+ to isolate the neural plate (NP) and the neural plate border (NPB). All the tissues were lysed and bisulfite converted with the EpiTect Plus Bisulfite Conversion Kit (Qiagen) following the manufacturer’s instructions. The regulatory regions of miR-137 were amplified by using sets of primers (see Table S2) from the bisulfite-converted DNA. The obtained products were gel purified and cloned into the pGEM-T Easy Vector (Promega). Individual clones were sequenced and analyzed.

## STATEMENTS AND DECLARATIONS

## Acknowledgments

We thank all the members of LBD for their contributions and helpful discussions during the course of our study.

## Authors’ Contribution

LAS, NV, and PHS-M designed and performed the experiments, analyzed the results, and wrote the paper.

